# Timing of Cyclic Estradiol Treatment Differentially Affects Cognition in Aged Female Rhesus Monkeys

**DOI:** 10.1101/248963

**Authors:** Mark G. Baxter, Anthony C. Santistevan, Eliza Bliss-Moreau, John H. Morrison

## Abstract

Some evidence suggests that there may be a limited "window of opportunity" for beneficial effects of hormone therapy on physiology after menopause in women. We tested, in aged, surgically menopausal (ovariectomized) rhesus monkeys, whether the timing of cyclic estradiol (E2) treatment impacted its effect on cognitive function. Monkeys were assigned to one of four treatment conditions after ovariectomy: either vehicle or E2 treatment for the duration of the protocol, vehicle for the first 2 years of the protocol followed by E2 for the remainder (delayed treatment), or E2 for the first year of the protocol followed by vehicle for the remainder (withdrawn treatment). Delayed treatment addressed the hypothesis that E2 treatment initiated more than 2 years after ovariectomy would have a reduced effect on cognitive function. Withdrawn treatment mirrors current clinical advice to women to use hormone therapy in the initial post-menopausal period then discontinue it. Two periods of cognitive testing assessed treatment effects on cognition over time. E2 treatment predominantly affected a prefrontal cortex-dependent test of spatiotemporal working memory (delayed response). Monkeys with delayed E2 treatment improved in delayed response performance over time, whereas vehicle-treated monkeys declined. Monkeys with withdrawn E2 treatment maintained their performance across assessments, as did monkeys treated with E2 across the entire protocol. These findings suggest that a "window of opportunity" for hormone treatment after cessation of ovarian function, if present in nonhuman primates, lasts longer than 2 years. It also supports the notion that beneficial effects of hormone therapy may persist after discontinuation of treatment.

---

The Women's Health Initiative (WHI) Memory Study (WHIMS) found that treatment of older women, aged 65 to 79 at study onset, with conjugated equine estrogen (CEE) alone, or CEE plus progestin (medroxyprogesterone acetate, MPA), had no beneficial effects on global cognitive function, was associated with decline in global cognitive function in some women, and increased the risk of mild cognitive impairment (MCI) and Alzheimer's disease (Espeland et al., 2004; S. R. Rapp et al., 2003b; Shumaker et al., 2004; 2003). These studies contradicted research conducted in animal models, as well as with human female participants, which supported the notion that hormone therapy (HT) after menopause benefits cognitive function (Hao et al., 2007; Markowska & Savonenko, 2002; P. R. Rapp, Morrison, & Roberts, 2003a; Voytko, 2000; 2002).

Considerable discussion and subsequent research has been devoted to understanding the findings from the WHIMS and analyzing its limitations (Craig, Maki, & Murphy, 2005; Maki, 2004; Sherwin, 2006; Sherwin & Henry, 2008). One possibility is that, women enrolled in the study may have been too far from the onset of menopause to benefit from HT. This hypothesis is consistent with the idea that there may be a critical period or "window of opportunity" for HT to have beneficial effects on cognitive function, such that benefits are seen when HT is initiated soon after menopause but not when it is delayed (Maki, 2006a; 2006b; Resnick & Henderson, 2002; Sherwin, 2006; Zandi et al., 2002). Alternatively, observational studies that identified beneficial effects of HT may have suffered from a "healthy user" bias, such that women that elect HT are in better health and better educated than women who do not.

Studies in animal models directly address many of the challenges present in studies of humans. Specific advantages include the ability to more precisely match treatment and control groups for chronological age, health status, life history, and other subject characteristics. Furthermore, direct control of formulation of HT, verification of compliance with treatment, exclusion of confounding factors such as diet and other medications that could modulate HT effects, and verification of treatment effectiveness in terms of circulating hormone levels are all possible in animal studies but difficult or impossible to achieve in observational or epidemiological studies with women. Nonhuman primates, specifically rhesus monkeys (*Macaca mulatta*), are well-suited for modeling the relationship between neuroendocrine and cognitive aging in humans, because many aspects of reproductive physiology are similar in rhesus monkeys and women (Gilardi, Shideler, Valverde, Roberts, & Lasley, 1997), and the profile of cognitive aging in rhesus monkeys parallels that of humans (Baxter, 2001; Herndon, Moss, Rosene, & Killiany, 1997; J. A. Roberts, Gilardi, Lasley, & Rapp, 1997). As observed in humans, prefrontal cortex dysfunction is a key signature, exemplified by impaired performance in the spatial delayed response task, indicating poor spatial working memory (Bartus, Fleming, & Johnson, 1978; Lyons-Warren, Lillie, & Hershey, 2004).

Although the outcome of the WHIMS has made it clear that HT is not suitable for preventing dementia or maintaining cognition in women over 65, many unanswered questions remain. In particular, the "window of opportunity" hypothesis (Resnick & Henderson, 2002; Sherwin, 2007) suggests that HT is only effective in maintaining cognitive function, and delaying the onset of dementia, if it is initiated within a limited time after menopause. On this view, women enrolled in the WHIMS were beyond the critical period, and thus did not benefit from HT. In contrast, in observational studies reporting beneficial neurocognitive effects of HT, women began treatment soon after the onset of menopause and so received the maximal benefit. This hypothesis finds support from some studies in animal models; aging rodents appear to experience maximal benefits from HT at a point in the lifespan that corresponds to late middle-age in humans (Frick, 2009). Spatial memory in rats is improved when HT is initiated immediately or 3 months after OVX, but not 10 months after OVX (Gibbs, 2000). Elevation of apical spine density in CA1 by estradiol (E2) is less effective 10 weeks post-OVX in rats with no intervening hormone treatment (McLaughlin, Bimonte-Nelson, Neisewander, & Conrad, 2008). Rhesus monkeys ovariectomized for 10 years or longer show no benefit from E2 treatment in their delayed response performance (Lacreuse, Wilson, & Herndon, 2002), although performance on a hippocampal-dependent spatial memory task is enhanced. Aside from the one study reporting an insensitivity of very long-term ovariectomized rhesus monkeys to E2, the question of the "window of opportunity" has received little attention in nonhuman primate models. Thus, we investigated in our model whether a substantial delay between surgical menopause and initiation of HT is associated with a reduction in the beneficial cognitive effects of HT.

A related question of considerable clinical significance is whether the beneficial effects of hormone therapy persist after discontinuation of treatment. In view of evidence from the WHI for increased risk of heart disease, stroke, blood clots, and breast cancer (Anderson et al., 2003; 2004; Wassertheil-Smoller et al., 2003), current clinical advice is for women to take hormone therapy for as short a period as possible after the onset of symptoms of menopause (for example, hot flashes), but to avoid chronic hormone therapy (Santen et al., 2010). Women randomly assigned to receive HT or placebo for 2-3 years immediately after menopause in a trial examining effects of HT on bone loss, showed lowered risk of cognitive impairment when examined 5-15 years later relative to women who had never taken HT, even if they discontinued HT after the end of the trial (Bagger et al., 2005). Testing whether beneficial effects of chronic HT persist in monkeys after discontinuation of treatment complements such studies, given the advantages of the monkey model outlined above, and the ability to determine whether corresponding changes in neurobiological markers also endure. Other therapies that improve cognition in aged animals, such as neurotrophic factors, result in improvements in cognition that persist after discontinuation of treatment (Frick, Price, Koliatsos, & Markowska, 1997). Evidence in favor of potential enduring benefits of HT after it is ended may inform women's choices about embarking on a short course of HT post-menopause, versus avoiding HT altogether.

To date, our studies in nonhuman primates have focused on a cyclic regimen of E2, which has beneficial effects on cognitive function in aged, surgically menopausal (ovariectomized) rhesus monkeys, as well as positive effects on density of dendritic spines and other indicators of "synaptic health" in prefrontal cortex (Hao et al., 2007; Hara et al., 2014; Morrison & Baxter, 2014; P. R. Rapp et al., 2003a). This contrasts with continuously administered E2 or E2 regimens combined with progesterone, which have sometimes not shown the same beneficial effects (Baxter et al., 2013; Ohm et al., 2012; but see Voytko,Murray, & Higgs, 2009; Kohama et al., 2016). Thus, in these studies, we used a cyclic E2 regimen (dosing E2 by injection every 21 days) similar to previous studies demonstrating positive effects of E2 on brain and cognitive function (Hao et al., 2007; P. R. Rapp et al., 2003a). To that end, monkeys in the different hormone conditions completed two standard tests of cognitive function widely used by our group - the delayed response task and the delayed nonmatching-to-sample task.

## Method

### Subjects and overview of experimental timeline

The experiments were performed at the California National Primate Research Center (CNPRC) under protocols approved by the University of California, Davis Institutional Animal Care and Use Committee. The initial subject group included 41 behaviorally-naive female rhesus monkeys (*Macaca mulatta*), aged 18.3-22.5 years at ovariectomy (age mean ± SD, 20.1 ± 1.1 years). Thirty-two of the monkeys were pair-housed with other monkeys on this study or colony monkeys, and 9 were singly housed. Pair-housed monkeys were separated at night to facilitate urine collection for hormone assays and monitoring of food intake.

Monkeys selected for inclusion in this study received physical examinations by a member of the CNPRC veterinary staff, to ascertain the presence of any health conditions that would confound results. Monkeys that passed the physical examination then received ovariectomy (OVX) surgery, a post-OVX evaluation of hormone status to verify OVX effectiveness, followed by assignment to treatment condition and beginning of one of four treatment protocols a mean of 109.5 days after OVX (range 63-175 days). As described in the introduction, there were 4 treatment groups: vehicle treatment throughout the experimental protocol (group Veh), E2 treatment throughout the experimental protocol (group E2), E2 treatment for the first ~11 months of the protocol (16 E2 treatments spanning 336 days) followed by vehicle treatment thereafter (group Withdrawn), or vehicle treatment for the first ~24 months of the protocol (35 vehicle treatments spanning 735 days) followed by E2 treatment thereafter until perfusion (group Delay). The initial number of monkeys in each condition was: Veh: N = 11; E2: N = 11; Withdrawn: N = 10; Delay: N = 9.

Our choice of these time intervals for the Withdrawn and Delay treatment groups were based in part on the 3:1 ratio of human:rhesus monkey in terms of lifespan (Tigges, Gordon, McClure, Hall, & Peters, 1988). An E2 treatment of about a year in duration in group Withdrawn would correspond to women using HT for about 3 years after the onset of menopause and then discontinuing it, consistent with many current clinical guidelines (Marjoribanks, Farquhar, Roberts, Lethaby, & Lee, 2017; Santen et al., 2010). For group Delay, E2 treatment would begin on average ~27 months post-OVX, allowing for about 3 months post-OVX washout and then 24 months of vehicle treatment; the actual range was 798-879 days (mean, 835.8 days). This corresponds to ~2.3 years post-OVX, or nearly 7 years in human lifespan. Although this does not equal the interval between menopause and start of HT in many of the WHIMS participants, it does represent a substantial period of time in the rhesus monkey lifespan without circulating ovarian hormones before HT is initiated, and one that was practical in terms of logistical constraints with working with aged monkeys.

Behavioral testing consisted of acclimation to the test apparatus, training to criterion and testing across delays on the delayed response (DR) task, and training to criterion and testing on the delayed nonmatching-to-sample task (DNMS). The first testing period began the day after the second E2 or vehicle treatment the monkey received, and the second testing period was timed to begin coincident with the day after the second E2 treatment in monkeys in the Delay group, 777 days after the beginning of the treatment protocol in each monkey. The first test period also incorporated DR and DNMS testing with distraction after completion of delay testing in each task, identical to that described in Baxter et al. (2013), but those results will not be presented here because distraction testing was not included in the second behavioral test period and our focus is on change in performance between the two test periods. Some monkeys also were tested in behavioral tasks in an automated, touchscreen-based apparatus (like that in Baxter et al., 2007) after completion of DNMS in the second behavioral test period but before perfusion, as part of pilot data collection for another planned study. These monkeys continued the hormone treatment they were receiving during the second behavioral test, either E2 or vehicle depending on their group assignment, until perfusion.

Nine of the 41 monkeys failed to complete the entire experimental protocol, with 5 going to perfusion early before the end of the first behavioral test period and another 4 between the end of the first behavioral test period and the end of the second. As a result, the final sample was N = 8 monkeys in each condition who completed the entire experimental protocol, with a time interval between OVX and perfusion averaging 3.35 years (range 2.8-4.0 years).

### OVX, washout, and hormone treatment

For OVX surgery, monkeys were sedated with 10 mg/kg ketamine i.m., given 0.04 mg/kg atropine s.c., then intubated, placed on isoflurane anesthesia to effect, and positioned in dorsal recumbency. A ventral caudal midline abdominal incision visualized the body of the uterus and both ovaries. Ovarian vessels and the Fallopian tubes were isolated, ligated, and severed. The abdominal wall was then closed in three layers with two layers of 2/0 absorbable suture and the final subcuticular layer with 3/0 absorbable suture. The animals were recovered in the CNPRC surgical recovery unit and given three days of post-operative analgesia, 1.5 mg/kg oxymorphone i.m. three times daily.

Urinary metabolites of estrogen (E1C) and progesterone (PdG) were analyzed by enzyme immunoassay in the Primate Assay Laboratory at the CNPRC as previously described by Shideler and colleagues (Shideler, Gee, Chen, & Lasley, 2001) to ensure intact ovarian activity before inclusion in the study and the success of the OVX surgery. For urine collection, cage pans were placed in the late afternoon for overnight sampling, and 3 ml of urine was collected by 0900 the following morning. Samples were collected daily for 6 weeks. Urine was centrifuged and decanted to remove any solid material and then frozen until analysis by enzyme immunoassay. Hormone concentrations were indexed to creatinine (Cr) to adjust for differences in urine concentration. These assays confirmed effectiveness of OVX, similar to the description in Baxter et al. (2013).

Monkeys receiving E2 treatment at any phase of the study received two i.m. injections of 100 μg estradiol cypionate in 1 ml peanut oil vehicle, 9 hours apart, with the first injection at approximately 0600 on the treatment day and the second at 1500. Pilot studies indicated this produced more consistent peak serum E2 levels after dosing compared to single E2 injections (as in Rapp et al., 2003). Monkeys receiving vehicle treatment received two i.m. injections of 1 ml peanut oil vehicle only on the same schedule. Treatments were given every 21 days.

### Behavioral testing

#### Delayed response(DR)testing

Testing took place in a manual test apparatus, identical to previous descriptions (O'Donnell, Rapp, & Hof, 1999; P. R. Rapp et al., 2003a; P. R. Rapp, Kansky, & Roberts, 1997). A white noise generator was used throughout training to mask extraneous sound. Acclimation and familiarization to the apparatus at the beginning of testing included offering the monkey food rewards in the test tray, as well as the opportunity to displace objects and plaques covering food wells in order to obtain reward. DR training was conducted in phases, with and without delays, as in previous studies (Baxter et al., 2013; P. R. Rapp et al., 2003a). Trials were initiated by raising the opaque barrier of the apparatus, and the monkey watched through a Plexiglas screen while one of the lateral wells of the stimulus tray was baited with a food reward (e.g., raisin or peanut). The lateral wells were subsequently covered with identical plaques and, during the initial phase of training, the clear barrier was raised immediately to permit a response (0 s delay). After the monkey displaced one of the plaques, the opaque barrier was lowered to impose a 20 sec intertrial interval (ITI). Daily test sessions consisted of 30 trials, with the left and right food wells baited equally often according to a pseudorandom sequence. Monkeys were trained until they achieve a criterion of 90% correct (9 errors or less in 9 consecutive blocks of 10 trials). Testing subsequently continued in an identical fashion except that a 1 sec delay was imposed between the baiting and response phase of each trial, and continued until criterion performance was re-achieved. In the next phase the memory demands of the DR task were made progressively more challenging by introducing delays of 5, 10, 15, 30 and 60 sec; testing was otherwise conducted as before (i.e. 30 trials/day, 20 sec ITI). Monkeys were tested for a total of 90 trials (3 days of 30 trials per day) at each retention interval.

#### Delayed nonmatching-to-sample (DNMS) testing

Testing took place in the same WGTA apparatus as DR. Trials in this task consisted of two phases, initiated when the opaque barrier of the WGTA was raised to reveal an object covering the baited central well of the stimulus tray. After the reward was retrieved, the opaque screen was lowered, and the sample item positioned over one of the lateral wells. The other lateral well was baited and covered with a novel object. During training, a 10 sec delay was imposed and recognition memory was tested by allowing monkeys to choose between the sample and the rewarded novel object. The discriminative stimuli were drawn from a pool of 800 objects according to a pre-determined sequence, ensuring that new pairs of objects were presented on every trial. Twenty trials per day were given using a 30 sec ITI, counterbalancing the left/right position of the novel items. Subjects were tested until they reached a 90% correct criterion by committing no more than 10 errors in 5 consecutive sessions (100 trials). The memory demands of the DNMS task were then made progressively more challenging by introducing successively longer retention intervals of 15, 30, 60, 120 sec (total = 100 trials each, 20/day), and 600 sec (total = 50 trials, 5/day). Monkeys remained in the test apparatus for all delays.

#### Statistical analysis

Acquisition (behavioral test period 1) or reacquisition (behavioral test period 2) of the DR and DNMS tasks in trials to criterion were compared between groups by either <u>t</u>-tests or oneway ANOVAs, depending on the number of conditions (2 or 4 in the first and second behavioral test periods, respectively). Mixed effects logistic regression was used to model the proportion of trials in which monkeys responded correctly on the DR and DNMS tasks during memory tests in which performance was challenged by progressively longer delays after criterion performance at short delays had been achieved. To account for repeated measures across time in monkeys, random intercepts were included for each monkey and random slopes were included for the time points on which monkeys were tested. Full three-way interactions between treatment group (Veh, E2, Delay, Withdrawn), time point (first or second behavioral test period), and delay interval on task (5-60 sec for DR and 15-600 sec for DNMS) were included in the DR and DNMS models in order to assess the degree to which treatment groups modified the change in performance on a given task over time relative to the vehicle control group, conditional on the difficulty of the task (i.e., delay interval between stimulus presentation and choice). Odds ratios (OR) between the odds of responding correctly on a given task at time follow-up time point relative to the initial time point represent measures of change in performance over time. The primary parameters of interest are the ratios between these ORs in experimental treatment groups (E2, Delay, Withdrawn) vs. the control group (Veh), which we report as ratios of odds ratios (ROR). A ROR significantly greater than 1 indicates that the change in performance over time either improved more, or declined less, in the indicated experimental group relative to Veh. Hypothesis tests were performed at the 0.05 level of significance and 95% confidence intervals are reported. All tests and confidence intervals were corrected using Dunnett’s method for comparing parameters from three separate experimental groups (E2, Delay, and Withdrawn) to the control group (Veh). All statistical analyses were conducted using R version 3.4.2 (R Core Team, 2014). We adopted this more sensitive approach for these analyses rather than, for example, repeated measures analysis of variance on percent correct scores across delay conditions, because it incorporates into the model random slopes for individual monkey variation, as well as the number of trials carried out at each delay. This increases the amount of information available about the odds of responding correctly on a particular trial, relative to a percent correct measure collapsed across trials.

## Results

### Delayed response

For initial acquisition during the first behavioral test, the Veh and Delay groups were collapsed into a single "vehicle" group and the E2 and Withdrawn groups into a single hormone treatment group, because at this point in the protocol they were being treated identically. During the first behavioral test, E2 treatment improved rate of acquisition of the DR task at the 0 sec, but not the 1 sec delay: trials to criterion at 0 sec, mean ± SD: vehicle, 267.8 ± 257.5, hormone treatment, 123.8 ± 168.4, *t*(39) = 2.13, *p* = 0.040; 1 sec, mean ± SD: vehicle, 344.7 ± 484.4, hormone treatment, 359.1 ± 500.8, *t*(39) = 0.09, *p* = 0.93. Reacquisition of the DR task at the beginning of the second behavioral test was not influenced by hormone treatment group (Veh, E2, Delay, Withdrawn), *F*(3, 28) < 1, *p* > 0.60.

Baseline performance on the DR task differed between at least two treatment groups on the 10 sec delay, *X*^2^(3) = 11.26, *p* = 0.01. Specifically, the odds of responding correctly with a 10 sec delay in the Delay group were half that of the Veh group (OR = 0.55, 95% CI [0.32 – 0.93], *p* = 0.03). Performance on the 10 sec delay task did not differ significantly between E2 and Veh (OR = 0.97, 95% CI [0.58 – 1.63], *p* = 0.99) or Withdrawn and Veh (OR = 0.62, 95% CI [0.37 – 1.05], *p* = 0.084). Furthermore, baseline performance on the DR task did not differ significantly between any two groups for the non-0 sec delays: 5 sec, *X*^2^(3) = 2.84, *p* = 0.42; 15 sec, *X*^2^(3) = 6.77, *p* = 0.08; 30 sec, *X*^2^(3) = 1.52, *p* = 0.68; or 60 sec, *X*^2^(3) = 0.537, *p* = 0.91.

In the memory test phase of DR in which performance was challenged with increasing delays, performance improved between the first and second test periods in group Delay compared to group Veh at short and intermediate delays, indicating effectiveness in the delayed E2 treatment begun before the second behavioral test in improving DR performance. With the exception of improved performance at the 15 sec delay in group E2 relative to group Veh between the first and second test periods, performance tended to be stable in the other treatment groups. Average performance at each time point is shown in Figure 1a. Average changes in performance across time are shown in Figure 1b. Figure 1c shows the differences between treatment groups in their change in performance over time. The change in performance over time in the 5 sec delay condition was 1.99 times (95% CI [1.04 – 3.82], *p* = 0.036) greater in group Delay relative to the change over time in group Veh. Change in performance over time on the DR task with a 5 sec delay did not differ significantly between groups E2 and Veh (ROR = 1.11, 95% CI [0.59 – 2.11], *p* = 0.94) or between groups Withdrawn and Veh groups (ROR = 0.86, 95% CI [0.46 – 1.61], *p* = 0.86). The change in performance over time in the 10 sec delay condition was 2.85 times (95% CI [1.53 – 5.30], *p* < 0.001) greater in group Delay relative to the change over time in group Veh. Change in performance over time on the delayed response task with a 10 sec delay did not differ significantly between groups E2 and Veh (ROR = 1.07, 95% CI [0.58 – 1.96], *p* = 0.97) or between groups Withdrawn and Veh groups (ROR =1.63, 95% CI [0.89 – 2.99], *p* = 0.15). The change in performance over time in the 15 sec delay condition was greater in groups Delay (ROR = 2.40, 95% CI [1.33 – 4.34], *p* = 0.001) and E2 (ROR = 2.14, 95% CI [1.18 – 3.89], *p* = 0.007) relative to group Veh. Change in performance over time did not differ significantly between groups Withdrawn and Veh (ROR =1.60, 95% CI [0.89 – 2.90], *p* = 0.150) at the 15 sec delay. There were no significant difference between any of the treatment groups and group Veh in the DR task at the 30 sec delay (Delay vs Veh: ROR = 1.37, 95% CI [0.77 – 2.43], *p* = 0.43; E2 vs Veh: ROR = 1.29, 95% CI [0.73 – 2.27], *p* = 0.57; Withdrawn vs Veh: ROR = 1.27, 95% CI [0.72 – 2.25], *p* = 0.62) or at the 60 sec delay (Delay vs Veh: ROR = 0.82, 95% CI [0.47 – 1.44], *p* = 0.72; E2 vs Veh: ROR = 1.01, 95% CI [0.58 – 1.76], *p* =1.00; Withdrawn vs Veh: ROR = 0.98, 95% CI [0.56 – 1.72], *p* = 1.00).

**Figure 1.**
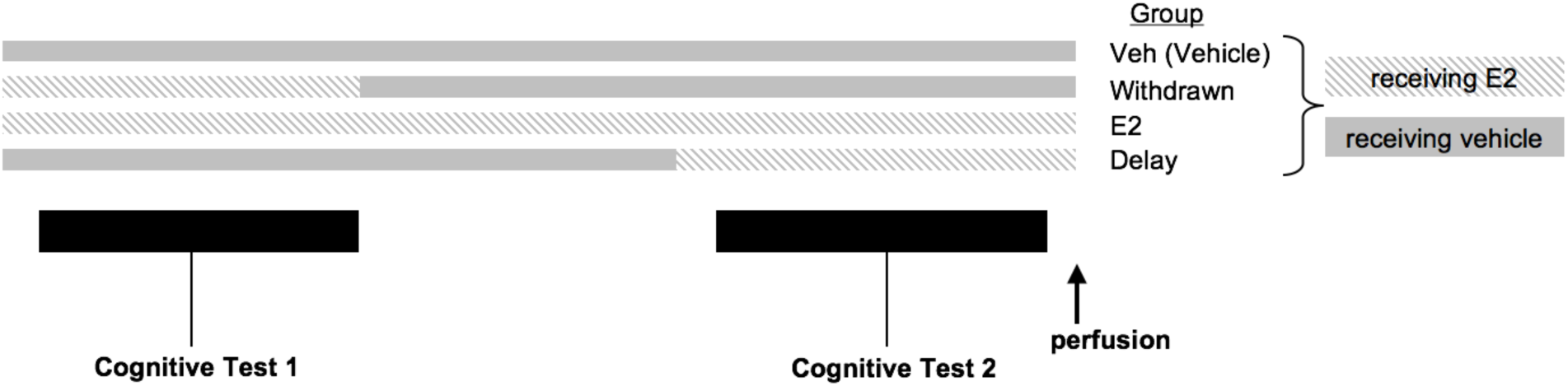
Schematic of the experimental timeline of hormone treatment and experimental groups. Group Veh (vehicle) received vehicle throughout the experiment, group E2 receives E2 treatment throughout the experiment, group Withdrawn receives E2 treatment for ~11 months and then begins vehicle treatment, and group Delay receives vehicle treatment for ~24 months and then begins E2 treatment. The two periods of cognitive testing are timed to coincide with E2 treatment in group Withdrawn, and then the onset of E2 treatment in group Delay.

### Delayed non-match to sample

As was the case for DR, for initial acquisition of DNMS during the first behavioral test, we collapsed experimental groups into two and compared "vehicle" and "hormone treatment" groups. E2 treatment did not affect rate of DNMS acquisition, trials to criterion at 10 sec mean ± SD: vehicle, 710,4 ± 620.8; hormone treatment, 812.7 ± 554.0; *t*(37) = 0.54, *p* = 0.59. Reacquisition of the DNMS task at the beginning of the second behavioral test was better in monkeys that were currently receiving or had previously received E2 (groups E2, Delay, Withdrawn) compared to monkeys in group Veh that had never received E2, means ± SDs:Con, 79.5 ± 117.5; Delay, 7.5 ± 14.9; E2, 0 ± 0; Withdrawn, 12.5 ± 23.75; *F*(3, 28) = 2.97, *p* = 0.049.

Baseline performance did not differ significantly between any two groups for any duration of delay (15 sec, *X*^2^(3) = 4.14, *p* = 0.25; 30 sec, *X*^2^(3) = 1.66, *p* = 0.65; 60 sec, *X*^2^(3) = 1.60,*p* = 0.66; 120 sec, *X*^2^(3) = 1.40, *p* = 0.71; or 600 sec, *X*^2^(3) = 1.31, *p* = 0.73).

Across all delays, change in performance between the first and second behavioral tests in the DNMS task did not differ between the three treatment groups (E2, Delay, Withdrawn) relative to the Veh group. Average performance at each time point is shown in Figure 2a. Average changes in performance across time are shown in Figure 2b. Figure 2c shows the differences between treatment groups in their performance over time. There were no significant differences between any of experimental groups (E2, Delay, Withdrawn) relative to group Veh at any of the delays tested in the DNMS task: 15 sec (Delay vs Veh: ROR = 0.74, 95% CI [0.37 – 1.48], *p* = 0.60; E2 vs Veh: ROR = 1.14, 95% CI [0.56 – 2.32], *p* = 0.92; Withdrawn vs Veh:ROR = 1.56, 95% CI [0.77 – 3.17], *p* = 0.32), 30 sec (Delay vs Veh: ROR = 0.76, 95% CI [0.38–1.49], *p* = 0.63; E2 vs Veh: ROR = 0.88, 95% CI [0.45 – 1.74], *p* = 0.92; Withdrawn vs Veh: ROR = 1.06, 95% CI [0.54 – 2.10], *p* = 0.98), 60 sec (Delay vs Veh: ROR = 0.72, 95% CI [0.38–1.36], *p* = 0.47; E2 vs Veh: ROR = 1.37, 95% CI [0.71 – 2.63], *p* = 0.52; Withdrawn vs Veh: ROR = 1.03, 95% CI [0.54 – 1.97], *p* = 1.00), 120 sec (Delay vs Veh: ROR = 0.91, 95% CI [0.49–1.69], *p* = 0.95; E2 vs Veh: ROR = 0.96, 95% CI [0.52 – 1.76], *p* = 0.99; Withdrawn vs Veh: ROR = 1.10, 95% CI [0.60 – 2.03], *p* = 0.94), or 600 sec (Delay vs Veh: ROR = 0.76, 95% CI [0.39 – 1.48], *p* = 0.63; E2 vs Veh: ROR = 0.91, 95% CI [0.47 – 1.78], *p* = 0.96; Withdrawn vs Veh: ROR = 0.99, 95% CI [0.51 – 1.94], *p* = 1.00).

**Figure 2.**
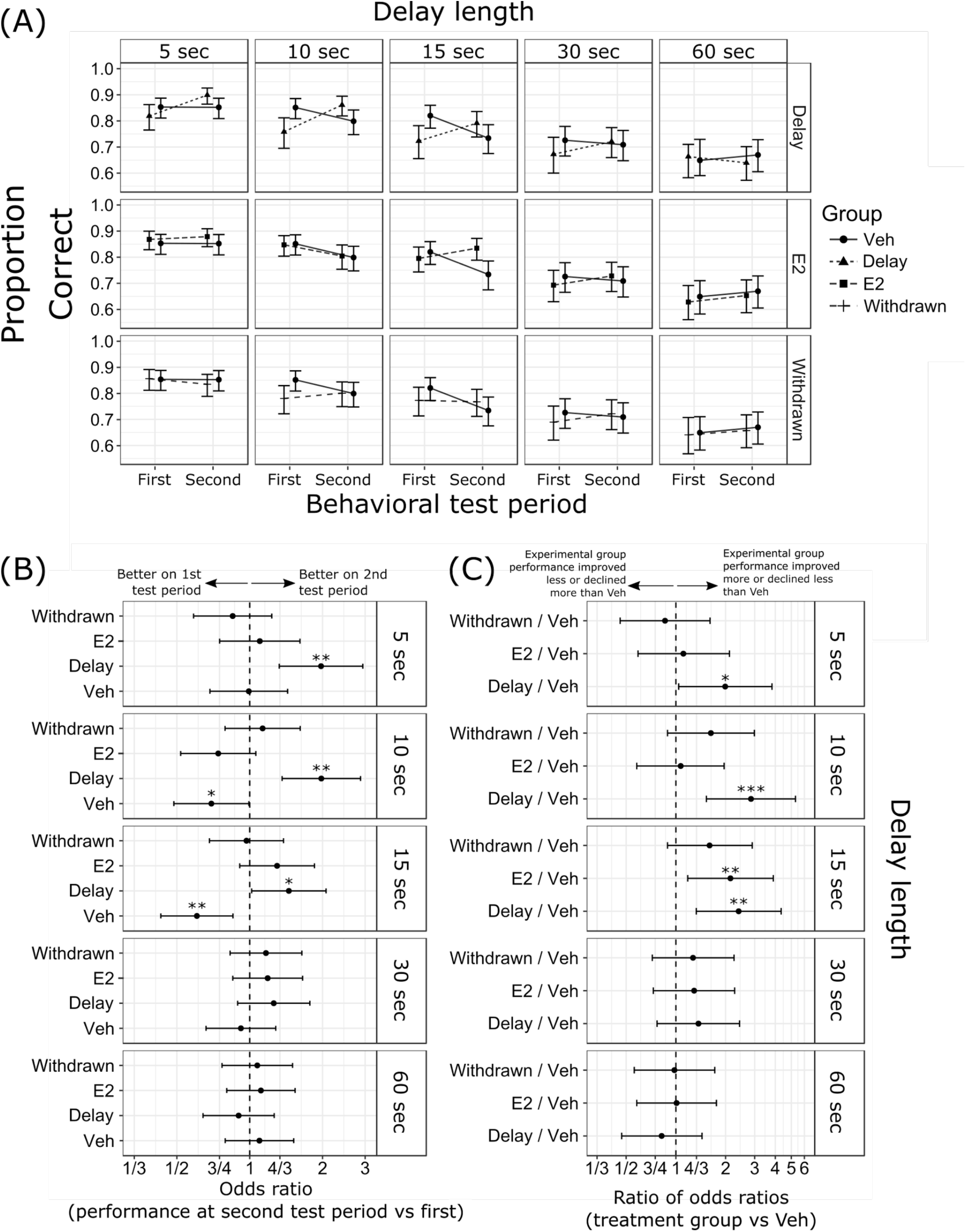
(A) Proportion of correct responses with 95% confidence intervals. (B) Odds ratios (OR) between performance at time second test period vs first test period, and (C) ratios of odds ratios (ROR) in performance across time in treatment groups relative to control. Results are separated by delay length (5 sec, 10 sec, 15 sec, 30 sec, and 60 sec delays). Group designations (Veh, E2, Delay, Withdrawn) as in Figure 1.

**Figure 3.**
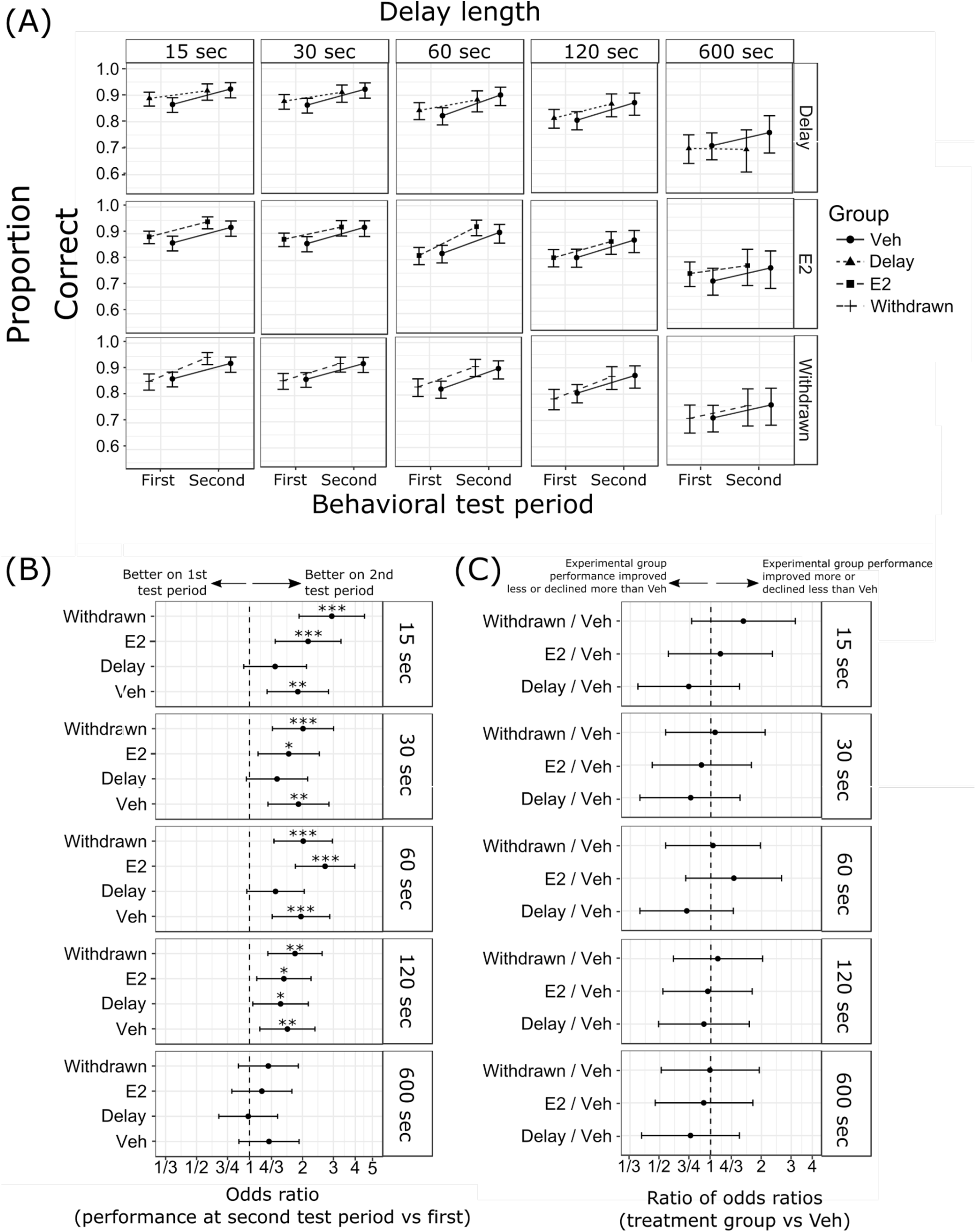
(A) Proportion of correct responses with 95% confidence intervals. (B) Odds ratios (OR) between performance at time second test period vs first test period, and (C) ratios of odds ratios (ROR) in performance across time in treatment groups relative to control. Results are separated by delay length (15 sec, 30 sec, 60 sec, 120 sec, and 600 sec delays). Group designations (Veh, E2, Delay, Withdrawn) as in Figure 1.

## Discussion

Our goals in this study were twofold. The first was to test whether E2 treatment, delayed by more than 2 years post-OVX, would still be effective in improving behavior in aged, surgically menopausal rhesus monkeys. The "window of opportunity" hypothesis derived from women in the WHIMS study would predict that E2 treatment after a substantial interval without circulating ovarian hormones would be ineffective. We found that prefrontal cortex-dependent spatial working memory in the DR task improved relative to baseline in monkeys with delayed E2 treatment, suggesting that if a "window of opportunity" exists in rhesus monkeys, it is longer than 2.3 years, the equivalent of ~7 years in humans. Whether comparison of time intervals in terms of lifespan ratio between monkeys and humans is valid in this regard remains an open question. That monkeys eventually become insensitive to effects of hormone therapy is suggested by one study where monkeys more than 10 years post-OVX did not experience any benefit of E2 treatment on the DR task (Lacreuse et al., 2002), but it is unclear whether this relates to the length of the post-OVX interval, the chronological age of the monkeys, or both. At the very least, our findings suggest that women who do not elect to begin HT immediately when menopausal symptoms are first experienced may reap some benefit from it. It is notable that there remain very limited data about the impact of HT in women that begin it in their 50s, around the average age of menopause (Marjoribanks et al., 2017).

Our second goal was to determine whether E2 treatment begun soon post-OVX and then withdrawn would produce any lasting beneficial effects on behavior. Memory performance in DR was stable in the second behavioral test relative to the first, indicating that performance did not significantly decline after E2 treatment was withdrawn, and monkeys in this group reacquired the DNMS task more readily than monkeys that had never received E2 post-OVX. This suggests that beneficial effects of E2 on cognition may persist after discontinuation of E2 in monkeys, providing a setting in which to investigate the cellular and molecular mechanisms of this effect.

The effects of hormone treatment were seen in the DR task, in initial acquisition where E2 improved learning of the task, in the pattern of change in delay performance between the first and second tests of DR, and in the reacquisition of DNMS in the second behavioral test. All of these effects suggest a locus of hormone treatment action in the prefrontal cortex, because both DR performance and DNMS acquisition depend on the integrity of the prefrontal cortex (Bachevalier & Mishkin, 1986; Goldman & Rosvold, 1970; Goldman, Rosvold, Vest, & Galkin, 1971; Shamy et al., 2011).

A limitation of our study is that across both types of testing, there was essentially no effect of E2 during baseline testing. This suggests that post-OVX interval used in this study, approximately 3.5 months, which was shorter than in previous studies in our group (~8 months in Rapp et al., 2003) may not be sufficiently long for synaptic health (Morrison & Baxter, 2014) and behavior regulated by impacted brain areas to be affected. Importantly, this did not compromise our ability to detect changes in behavior over time. However, this suggests a period of resilience of behavior following loss of circulating ovarian hormones before synaptic health deteriorates to a point that cognitive impairments are evident. This finding also gives an indication of the importance of timing of HT and is an idea that our future research will continue to investigate.

## Author Note

Mark G. Baxter, Department of Neuroscience and Friedman Brain Institute, Icahn School of Medicine at Mount Sinai; Anthony C. Santistevan and Eliza Bliss-Moreau, Department of Psychology, University of California, Davis, and California National Primate Research Center; John H. Morrison, Department of Neurology, University of California, Davis School of Medicine, and California National Primate Research Center.

This project was supported by National Institute on Aging (NIA) award P01-AG016765. The California National Primate Research Center is supported by National Institutes of Health (NIH) Office of the Director award P51-OD011107. The content is solely the responsibility of the authors and does not necessarily represent the official views of the NIH. We thank Mary Roberts, Tracy Ojakangas, and Lisa Novik for technical assistance, and Nancy Gee and Bill Lasley for hormone assay data verifying ovariectomy.

Correspondence concerning this article may be addressed to Mark Baxter, Department of Neuroscience, Icahn School of Medicine at Mount Sinai, One Gustave L. Levy Place Box 1639, New York, NY 10029.

